# The palisade-specific gene *IQD22* enhances photosynthetic capacity by phenocopying “sun leaf” architecture

**DOI:** 10.1101/2022.08.27.505449

**Authors:** Michaela C. Matthes, Joao P. Pennacchi, Johnathan A. Napier, Smita Kurup

## Abstract

The world requires a rise in crop production which needs to be increased significantly in order to satisfy demand by 2050^1^. However traditional plant breeding approaches are anticipated to fall short in delivering the increases in yield required and therefore targeted manipulation of plant metabolism is increasingly being pursued with the aim to achieve this goal^2^. Improving photosynthetic efficiency is predicted to have a significant influence on enhancing crop productivity and several ambitious genetic engineering projects are currently under way to achieve higher photosynthetic rates in crops^3,4^. A naturally evolved adaptive trait which allows plants to increase photosynthetic efficiency specifically under high light is the differentiation of ‘sun’ leaves which are characterised by an increase in palisade cell layers and more elongated cells within this layer^5^. These morphological changes allow a more efficient distribution of light within the leaf and provide an increased cell surface to which chloroplasts can relocate thereby increasing the capacity for CO_2_ up-take^6^. Here we show that, surprisingly, this complex morphological trait can be phenocopied by the modulation of the expression of a single palisade specific gene, *IQD22*, in *Arabidopsis*. Furthermore, we could show that the architectural changes were reflected in an increase of photosynthetic rate of 30%. The simplicity with which we could enhance photosynthesis by phenocopying sun leave traits is in stark contrast to the complex and challenging metabolic engineering approaches currently being pursued.

---

Meeting the food demands for the growing human population will require a step change in improving current crop productivity as the predicted trajectory for crop yields is insufficient to feed the world’s population by 2050^1^. Besides increasing resilience of crops to biotic and abiotic stresses, enhancing photosynthetic efficiency is considered to also be a major contributor to the improvement of yield and an increase in biomass^2-4^. So far, the majority of approaches to optimise photosynthetic efficiency has been aimed at engineering and altering enzyme activities involved in metabolic pathways linked to the photosynthetic process such as enhancing the efficiency of the Calvin-Benson cycle, reducing energy loss through photorespiration and acceleration of the recovery from photoprotection (non photochemical quenching (NPQ))^7-9^. More recently the successful introduction of a simplified cyanobacterial CO_2_-concentrating mechanism (CCM) into tobacco chloroplasts has been reported which suggests that an increase in CO_2_-fixation, reduction of water loss and lower investment in Rubisco through reconstitution of a functional carboxysome in C3 plants might ultimately be feasible^10^. In an alternative approach Hart et al., 2019 engineered the photocycles of phototropin 1 and 2, two blue light photoreceptors which play important roles in regulating several responses that optimise photosynthetic efficiency, and could show that this lead to improved biomass accumulation specifically under light-limiting conditions^11^. Probably the most ambitious approach envisages the conversion of C3 crops into C4 plants as this requires the introduction of both, leaf architectural and physiological traits. These include the differentiation of photosynthetically active vascular bundle sheath cells, modification in the biochemistry of several enzymes and the modification of the inter- and intracellular transport of metabolites^12^. The transfer of such a complex trait system into C3 plant in one single step is therefore highly unlikely.

Besides the evolution of C4 photosynthesis, another naturally evolved trait in plants to maximise photosynthetic efficiency, specifically under high light conditions, is the development of ‘sun’ leaves which are characterised by a palisade mesophyll that comprises more cell layers and more elongated cells than their shaded counterparts^13,14^. The columnar shape of palisade cells has been functionally linked to a more efficient distribution of light throughout the mesophyll and their increased cell height facilitates a better distribution of chloroplasts along the anticlinal cell walls which enhances CO_2_ uptake and improves photosynthesis^15-17^. This raises the question whether uncoupling increases in leaf thickness from its environmental adaptive response and targeting it for genetic manipulation could lead to photosynthetically more efficient plants but recent attempts to elucidate the genetic mechanisms underlying the development of thick leaves suggested that this trait has a complex genetic basis and, similar to the C4 trait may be recalcitrant to engineering^18,19^. However, here we report that surprisingly the expression of a single gene can confer the anatomical and physiological traits of sun leaves to *Arabidopsis thaliana* leaves.

*Arabidopsis* leaves generally have poorly-developed palisade cells when grown under standard conditions (60-250 μ ml m^-2^ sec^-1^ white light), however under higher fluence rates (360 – 600 μ ml m^-2^ sec^-1^ white light) leaves were shown to be able to acclimate by a moderate increase of the anticlinal elongation of the palisade mesophyll cells and the differentiation of a second layer of palisade cells^20,21^. We fortuitously discovered that expression of IQD22 (At4g23060) translationally fused 3’ to mCherry and driven by its native promoter, generated T1 plants which developed leaves which were rounder and much thicker than those of untransformed WT plants, whereas other parts of the plant showed no phenotypic changes (Fig 1a and Supplemental Fig S1). IQD22 is a member of the plant-specific IQ67 DOMAIN (IQD) family of proteins which is defined by a characteristic domain of 67 conserved amino acid residues containing two to three repeats of the IQ motif (IQxxxRGxxxR) (Supplemental Fig S2)^22^. So far, the function of this large family (33 members in *Arabidopsis*) has remained elusive with a possible role reported for IQD1 in the regulation of glucosinolate metabolism (Arabidopsis) and for IQD12 in the regulation of fruit shape in tomato^23,24^. More recently it was shown that many members of the IQD67 family localise to plasma membrane subdomains, nuclear compartments and frequently the microtubule array^25^. The IQ67 domain has been shown to be required and sufficient for interaction with calmodulin *in vitro* and it has been suggested that IQD proteins are involved in integrating Ca^2+^ -calmodulin signalling with microtubule associated processes such as regulation of cell function, shape and growth^26-28^. However, the biological functions of these IQD proteins are still largely unknown and their role in microtubule organisation remains uncharacterised. In order to better understand the noted increase in leaf thickness in the IQD22 expressing lines, we performed cryo-fracture on WT and transgenic leaves for analyses using scanning electron microscopy. As shown in Fig 1b, the leaf of the IQD22 expressing line displayed the typical traits of a ‘sun’ leaf with a well-developed palisade tissue containing more cell layers with more elongated cells when compared to WT, whereas the spongy mesophyll cell layer remained largely unaffected (Fig 1b). For an in depth anatomical characterisation of this phenotype we generated homozygous T3 lines and prepared resin-embedded cross sections of fully expanded leaves of 3-week old plants. Leaf thickness was increased by >50% in three independent transgenic lines when compared to WT (Fig 1c) which was mainly the result of changes in palisade tissue architecture. In WT, palisade tissue consisted of one to two cell layers whereas in the transgenic plants three and sometimes four cell layers could be observed (Supplementary Fig S3). To compare cell slenderness of the palisade cells, height-to-width ratios were calculated. A remarkable shift towards more slender and elongated palisade cells in palisade cell layer one and two in the lines expressing pIQD22:IQD22:mCherry (Fig 1d) was observed. A view of the paradermal plane of fully expanded leaves revealed that palisade cells were on average smaller and arranged more loosely generating wider intercellular spaces in the transgenic lines compared to WT (Supplementary Fig S4). In order to determine whether the increase in leaf thickness was dependent on the light intensity in which the plants were grown, we grew WT and transgenic lines under 60 μ mol m^-2^ s^-1^ white light for 7 weeks and sampled fully expanded leaves for anatomical characterisation. Leaf morphology was effectively unchanged between the different genotypes, however a slight increase of leaf thickness occurred in the pIQD22:IQD22:mCherry lines due to some elongated palisade cells (Supplementary Fig S5). This demonstrates that also in the pIQD22:IQD22:mCherry lines, leaf thickness is dependent on light fluence rate. In summary, increasing the dosage of IQD22 in *Arabidopsis* by introducing IQD22 under the control of its native promoter renders the palisade tissue more sensitive to light fluence so that it manifests the anatomical characteristics of ‘sun’ leaf palisade tissue at light levels where this normally would not occur.

**Fig 1:**
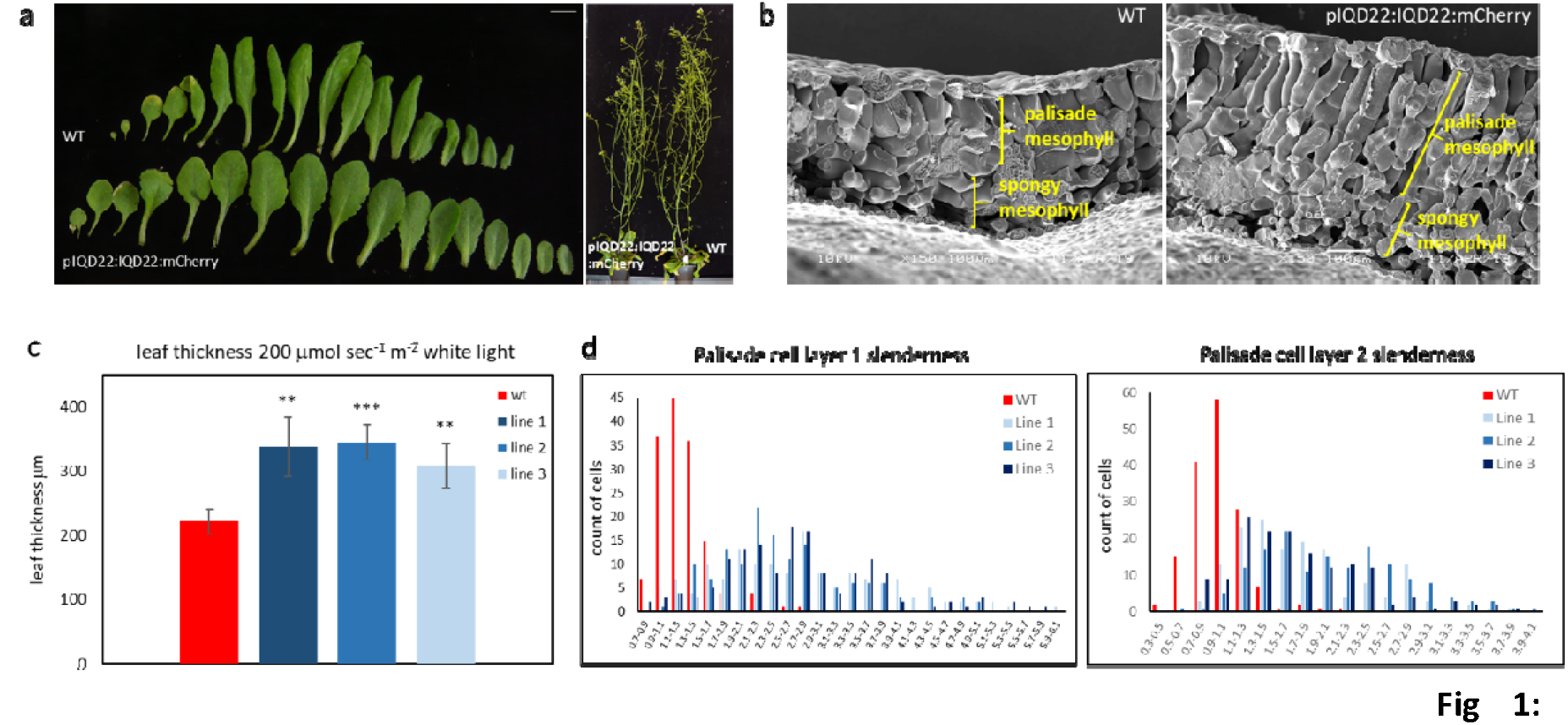
Phenotype and characteristics of Arabidopsis lines expressing IQD22 under the control of its native promoter. **a**, leaf series of 5 week old plants grown under long day conditions in high white light (200 μmol m^-2^ sec^-1^); fully mature flowering plants display altered morphology solely in leaves of transgenic plants (see also Fig S1). **b**, Cryo-fractured mature leaf of a 5 week old WT plant and a representative transgenic line expressing pIQD22:IQD22:mCherry. Plants were grown under high white light and long day conditions. Anisotropic cell elongation of palisade cells in the transgenic lines can be observed. Bar = 100μm. **c**, Leaf thickness of pIQD22:IQD22:mCherry lines is increased by ca 50%. Measurements were performed on images of resin embedded cross sections of leaves for three independent lines using ImageJ and are based on n=3 per line. **d**, Frequency distribution of palisade cell slenderness (length/width) as measured in ImageJ shows a shift towards more elongated cells in three transgenic lines when compared to WT; n= >150 cells from leaf sections of three individual plants per line.

The observation that palisade cells are the cells most responsive to the expression of the pIQD22:IQD22:mCherry transgene is explained by the fact that it is solely these cells which express IQD22 *in planta*. pIQD22:GUS reporter lines showed that IQD22 is expressed predominantly in leaves and cross sections of resin-embedded samples localised this expression specifically to palisade cells (Fig 2a and Supplementary Fig S6). Further confirmation of this finding was obtained by analysing the subcellular localisation of IQD22 by confocal microscopy of the *Arabidopsis* pIQD22:IQD22:mCherry lines. Analysis of the paradermal planes of the adaxial and abaxial sides of mature leaves by confocal microscopy detected no mCherry fluorescence in the adaxial or abaxial epidermis, however, localisation of the fusion protein suggested a possible co-localisation with the cortical microtubule array of the palisade mesophyll (Fig 2b). To unambiguously confirm co-localization of IQD22 with microtubules we generated transgenic *Arabidopsis* lines expressing the translational IQD22:mCherry fusion under the control of the constitutive 35S promoter. A line was crossed to *Arabidopsis* expressing α-tubulin fused to GFP and F1 plants were analysed for localisation of fluorescent signals. As shown in Fig 2c, in adaxial epidermal cells the signal emanating from α-tubulin-GFP overlaps perfectly with that generated by 35S:IQD22:mCherry corroborating a close association of these proteins *in vivo*. Since cortical microtubules play a pivotal part in controlling cell growth and shape, these findings suggest that it is through this interaction with microtubules in the palisade cells that increased expression of IQD22 is responsible for the observed altered morphology, however the underlying mechanisms await further investigation.

**Figure 2:**
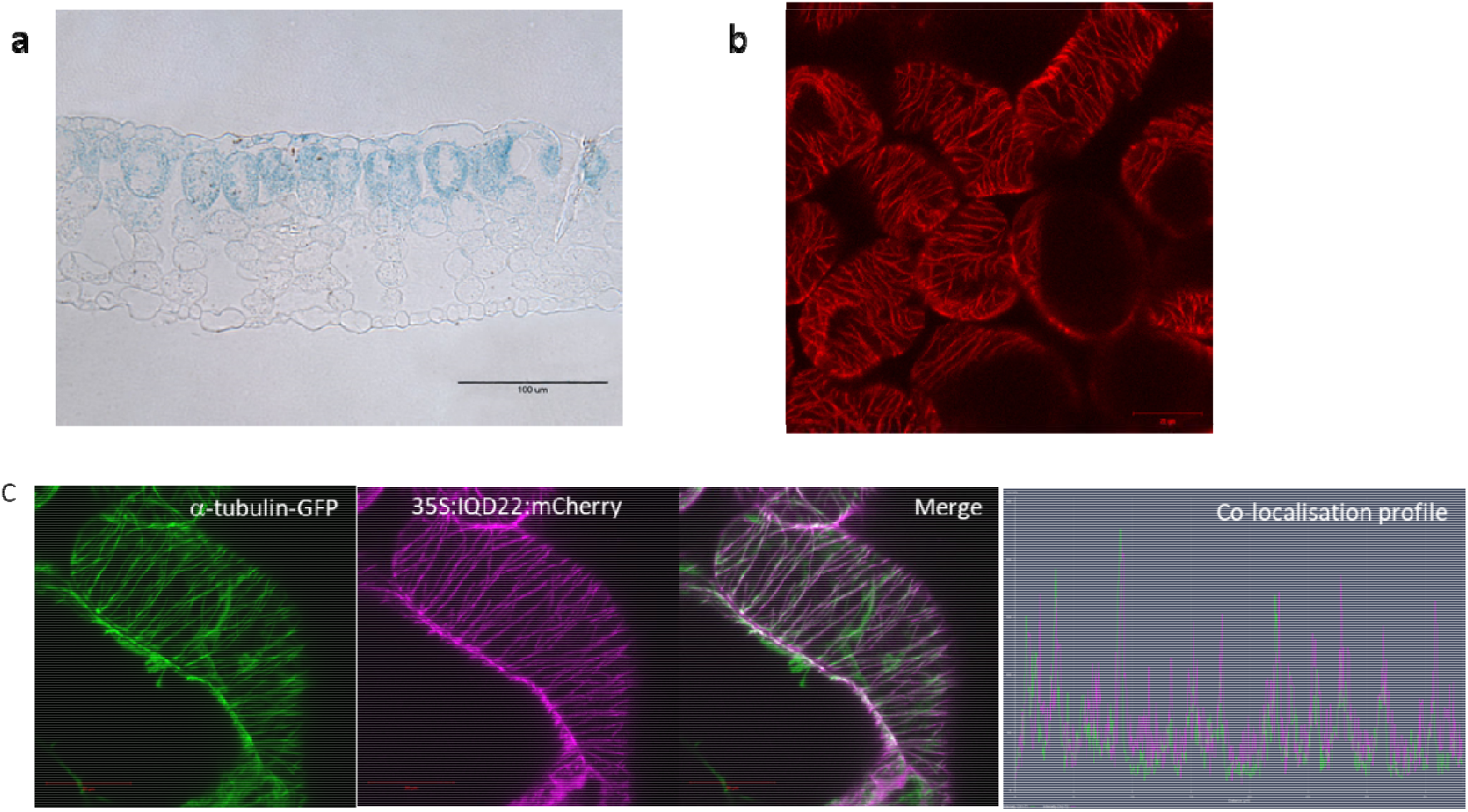
Cellular and sub-cellular localisation of IQD22 expression. **a**, IQD22 is expressed specifically in the palisade cells of Arabidopsis leaves. Leaves of transgenic plants expressing the pIQD22:GUS reporter construct were processed for histochemical localisation of GUS, fixed, embedded in resin and sectioned. A representative 4 μm section from one of three independent lines is displayed. Bar= 100 μm. **b**, Maximum intensity projection of a paradermal confocal image of the adaxial surface of a mature leaf of pIQD22:IQD22:mCherry line. This was a fully expanded leaf of a three week old plant grown under long day conditions under high white light. Bar = 20μm. **c**, co-localization of IQD22 with the cortical microtubule array. Image of an adaxial leaf epidermal cells from a F1 plant generated from a cross between *Arabidopsis* Col-0 α-tubulin-GFP and *Arabidopsis* Col-0 35S:IQD22:mCherry line. The co-localisation profiles shows overlap of the GFP and mCherry signal. Bar = 20μm.

Blue light perceived by phototropins plays a pivotal role in regulating several responses which contribute to the optimisation of photosynthetic efficiency, among them the development of cylindrical palisade cells which in *Arabidopsis* is controlled specifically by Phototropin 2^29,30^. We therefore asked the question if IQD22 could be involved in this process by analysing the response to blue light in the following genotypes: *Arabidopsis* Col-0 WT, two transgenic lines expressing pIQD22:IQD22:mCherry (line 1 and 4), a *phot2* KO line (Salk_124475) and a line homozygous for *phot2* and pIQD22:IQD22:mCherry. We had intended to include IQD22 KO plants in this analysis, however three independent SALK lines for this gene (SALK_110842, Salk_118021 and Salk_119835) which had been genotyped for homozygosity and which were confirmed by RT-PCR for absence of the transcript were subsequently shown by RNA-seq to be knock-downs (Supplementary Fig S7). Additionally, as IQD67 is a family with 33 members, redundancy may also pose a problem when interpreting data from mutants of individual genes. The genotypes were grown at 60 μmol m^-2^ s^-1^ white light (low light) for 20 days and then transferred to a LED cabinet with broad spectrum blue light at 200 μmol m^-2^ s^-1^ (high light) and grown under these conditions for an additional 2 weeks. Samples were taken of leaves 9 and 10 which had been just visible at the time of transfer into blue light conditions for morphological analyses by cross-sectioning. Figure 3a shows representative cross sections of leaf 10 of all the genotypes used. Expression of pIQD22:IQD22:mCherry led to up to a 60% increase in leaf thickness compared to WT, whereas no such increase was detected when this expression occurred in a *phot2* background (Fig 3b). This increase results from a combination of more palisade tissue layers and increased anticlinal elongation of the cells within this tissue. In the case of WT, palisade cells had acquired an elongated cell shape in blue light conditions which was however much more pronounced in transgenic lines with individual cells being up to 1.8 time longer than in WT (Fig 3c). This pronounced palisade cell elongation depends to a large extent on the functional blue light receptor phototropin 2 as in the *phot2* background it did either not occur or was much reduced with a maximal elongation similar to WT.

**Fig 3:**
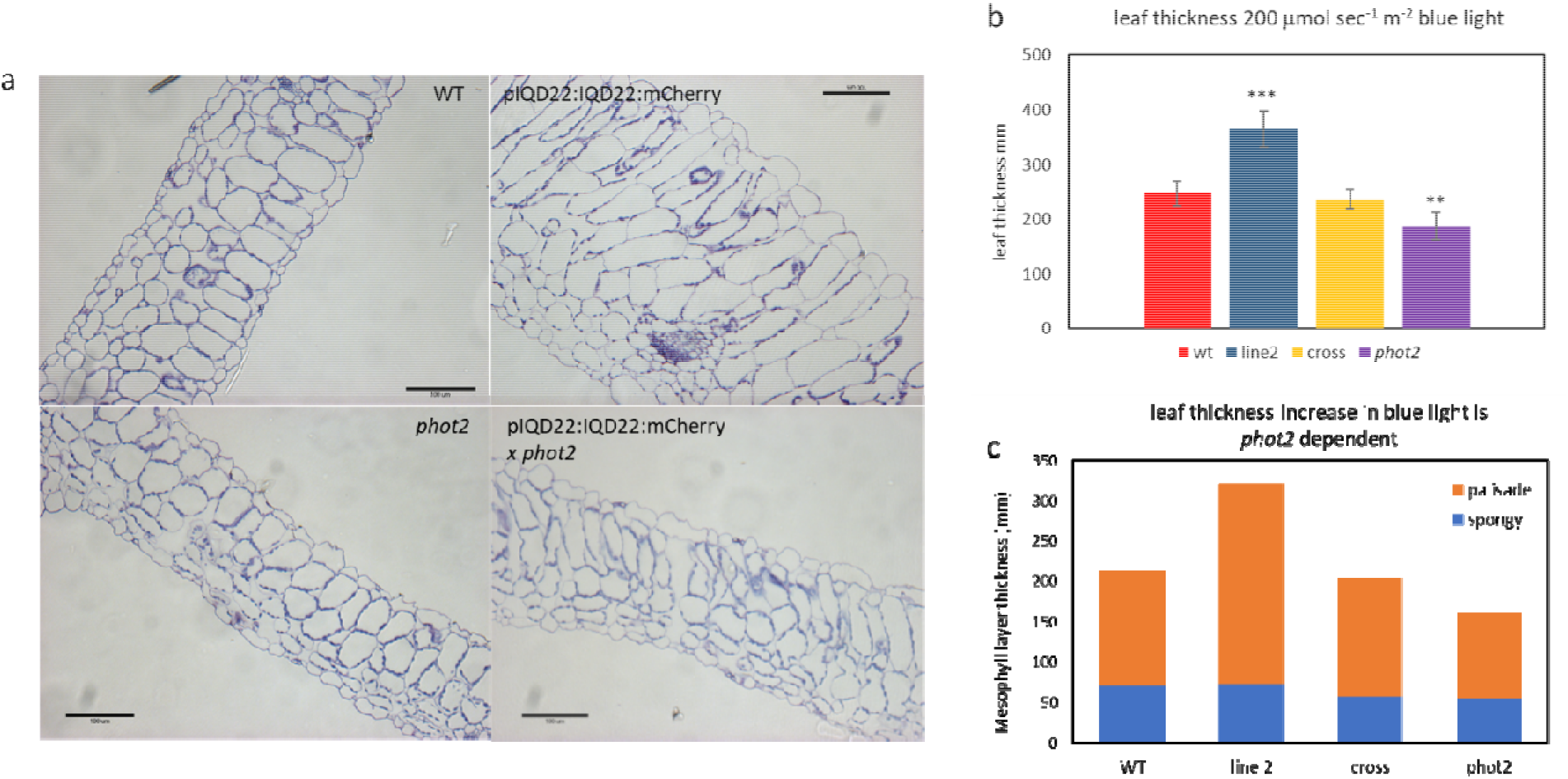
Palisade cell elongation in pIQD22:IQD22:mCherry lines is regulated by blue light. **a**, Cross sections, 2 μm in thickness, of resin embedded leaf samples of leaf 10 from WT, pIQD22:IQD22:mCherry line 2, *phot2* and pIQD22:IQD22:mCherry in a *phot2* background after transfer from low white light (60 μmol m^-2^ s^-1^) to high (200 μmol m^-2^ s^-1^) blue light for two weeks. Bar = 100 μm. **b**, Leaf thickness measurements in WT, pIQD22:IQD22:mCherry line 2, phot2 and pIQD22:IQD22:mCherry in a phot2 background. Measurements were performed in ImageJ and are based on n=3 per genotype or n=5 for the cross (pIQD22:IQD22:mCherry x phot2). Leaf thickness of Line 2 is significantly greater than WT (t-test, p<0.001), whereas it is similar to WT in the cross (phot2 background). However, the decrease in leaf thickness of the cross is less than that observed when *Phototropin2* is lacking. **c**, The combination of less cell layers in the palisade mesophyll and/or reduced elongation of those cells is the underlying cause that leaf thickness in the *phot2* background is diminished whereas the thickness of the spongy mesophyll layer remains largely constant in all the genotypes analysed.

Palisade cell morphology is intimately linked with an increase in photosynthesis in a high irradiance environment^5,6^. Since an increase in the expression of IQD22 in palisade cells of *Arabidopsis* leaves leads to the morphological differentiation of typical ‘sun leaves’ we next investigated whether this was also reflected in photosynthetic parameters. As shown in Figure 4a, we could measure a net increase in the photosynthetic rate of the transgenic lines compared to WT with a 30% higher capacity for fixing carbon per unit leaf area. Stomatal index between WT and transgenic lines however was similar whereas a slight but not significant increase of stomata per unit area was noted for IQD22 overexpressing lines (Supplementary Fig S8). Several parameters contribute to this increase in photosynthetic rate.

**Fig 4:**
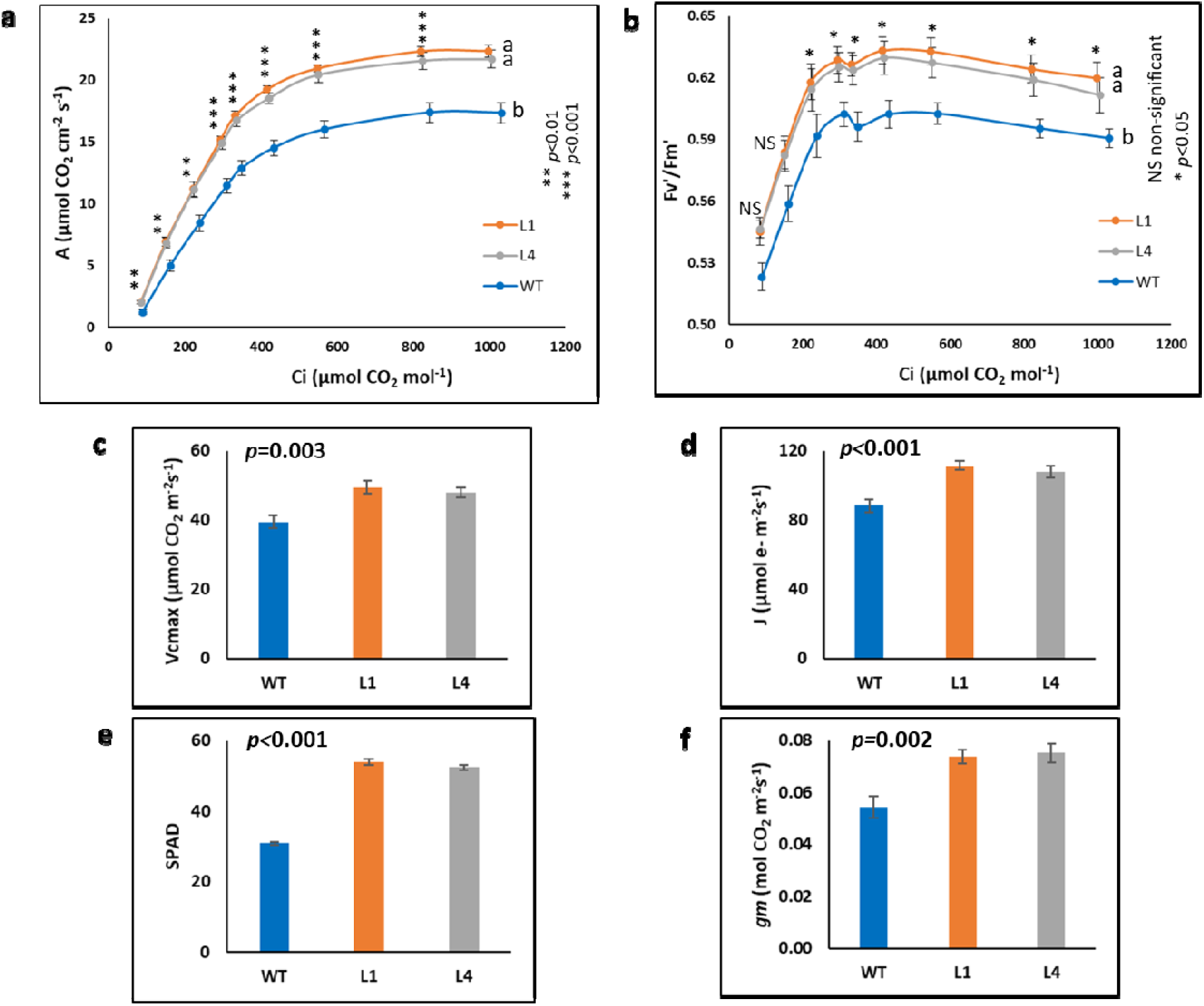
Measurements of photosynthetic parameters of pIQD22:IQD22:mCherry lines and WT. Photosynthetic measurements were performed on mature leaves of 5 week old WT and pIQD22:IQD22:mCherry lines 1 and 4 grown under high white light (n=5). **a**, response of net CO_2_ assimilation (A) to the intercellular CO_2_ concentration (Ci). **b**, response of light-adapted fluorescence (Fv’/Fm’) to the intercellular CO_2_ concentration (Ci). **c**, maximum carboxylation rate (Vcmax). **d**, electron transport rate (J). **e**, chlorophyll content (SPAD). **f**, total conductance between intercellular spaces and chloroplast (gm). Averages with different letters within the figure panel (a and b) presented significant differences at the indicated p level through the Tukey test. * in Figure panels a and b indicates significant differences (** p value1⍰<⍰10.01; *** p value1⍰<⍰10.001), judged by the Tukey test.

Mesophyll conductance, *gm*, is increased by 37% (Fig 4f) in the transgenic lines implying that CO_2_ delivery to the chloroplasts (and probably to Rubisco) is more efficient, probably due to a combination of the looser arrangement of the mesophyll cells and an increase of the chloroplast surfaces aligned along the columnar palisade cell wall. Compared to WT, chlorophyll content in the transgenic lines was increased by 52% as determined using a SPAD meter (Fig 4e) enabling the harvest of more light. An increase in electron transport rate (J; +24% Fig 4d) and efficiency of light use (Fv’/Fm’; +5% Fig 4b) by the components of photosystem II also contributes to the higher net photosynthetic rates. Finally, the capacity and velocity of the leaf to fixate carbon in the biochemical reaction through the Calvin-Benson cycle, displayed as the maximum velocity of carboxylation of Rubisco (Vmax) in Fig 4d, shows an increase by 23% in the transgenic lines. In summary, both the photochemical and biochemical steps of the photosynthetic process are enhanced in the transgenic plants.

This is the first report of a palisade specific protein, IQD22, that controls the anatomical and physiological characters of leaves in *Arabidopsis* by the modulation of its expression. Whether other species respond in a similar way needs to be determined, although preliminary results suggest that similar effects are obtained in *Camelina sativa* and poplar but how major crops will respond remains to be seen. With the recent rising interest in enhancing the efficiency of photosynthesis in order to achieve future increases of biomass and yield of our major crops, we propose that genetically modifying leaf architecture by manipulating IQD22 expression represents a powerful yet simple method. Furthermore, this morphological approach is distinct from previously described metabolism-focused efforts to boost photosynthesis by overexpression of enzymes involved in carbon assimilation, and represents a promising new avenue for further exploration.

## Experimental Methods

### Plant material

The Col-0 ecotype of *Arabidopsis thaliana* was used for the studies reported here. We thank Dr Joseph F. Mckenna (Oxford Brookes University) for the 35S-TUA6-GFP seed. The *phot2* KO Salk_142275 line was obtained from NASC. Unless otherwise stated, plants were grown in a Weiss Gallenkamp cabinet under long day conditions (16 h light 23°C/ 8 h dark 18°C) and 250⍰⍰mol m^-2^ sec^-1^ white light provided by a fluorescent lamp.

### Blue light treatment

Blue light treatment was performed according to López-Juez et al (2007) with some modifications. In brief, 7 days old seedlings were transferred into compost and grown in a MobyLux GroBanks cabinet from CLF Plant Climatics under 8 h light (23°C) and 16 h dark (18°C) and 60 1mol m^-2^ sec^-1^ white light provided by a fluorescent lamp for 2 weeks. After 2 weeks plants were transferred to a Weiss Fitotron cabinet fitted with a LED RX 30 lamp from Heliospectra. Blue light at 20011mol m^-2^ sec^-1^at 30 cm distance from the lamp, where plants were placed, was achieved using the 420 nm and 450 nm wavelength channels with the 450 nm channel set at maximum intensity (1000) and the 420 nm channel at intensity 410. Plants were grown in long day conditions for an additional 2 weeks. Sampled leaves were selected as the most recent leaf that was ca. 2 mm long (just visible) when the treatment started and that had fully expanded under the treatment irradiance. This corresponded to leaves 9 or 10 under conditions described here.

### Leaf sections

5 mm wide strips in the middle of the leaf were cut extending from the mid-vein to the leaf edge and were fixed in 4% paraformaldehyde and 2.5% glutaraldeyde in 0.05M phosphate buffer pH 7.2 for 1 h at room temperature and then at 4°C overnight. The fixed tissues were gradually dehydrated with ethanol and embedded in LRWhite Resin (Agarscientific). 1–2 1m sections were cut with a Reichert Ultramicrotome (Ultracut; Reichert-Jung, Vienna, Austria) and stained with toluidine blue.

### Microscopy

Light microscopy was performed using a Zeiss AxioPhot upright microscope and images were acquired using a Q-Imaging Retiga digital camera. For live cell imaging, fluorescence from GFP-or mCherry-tagged proteins was observed under a confocal laser scanning microscope (LSM780; Zeiss) with the following excitation and emission filters: GFP (488 and 525 nm) and mCherry (561 and 595 nm). Samples for cryoscanning electron microscopy were first frozen in slush, prepared in an Gatan Alto 2100 cryo-system (Gatan), and then analysed in a JEOL JSM-6360 LV SEM. Leaf cross sections were obtained by fracturing the leaf blade after freezing.

### Plasmid constructs

For the GUS reporter fusion, 1.6 kb of the IQD22 (At4g23060) promoter region was amplified with the primers SphI F 5’-GCATGCCAACTTGCATTATTACTGTACCA-3’ and NcoI R 5’-CCATGGCTAATGAAAGTTACTTGACGA-3’. The sequence for the NcoI R primer does not correspond to the sequence deposited in TAIR as we had determined that 19 bp directly adjacent to the 5’ side of the IQD22 ATG start codon differ from the TAIR sequence in the genotype we used for amplification. The amplified fragment was cloned into pJET1.2 using the cloneJET PCR cloning kit (ThermoFisher), excised with SphI and NcoI and cloned into pJD330 from which the S35 promoter had been excised by SphI and NcoI. The whole pIQD22:GUS fusion was amplified with primers containing AscI sites (AscI F 5’-GGCGCGCCCTTTGCCAACGAATGTTAGTTGT-3’ and AscI nos 5’-GGCGCGCCATCTAGTAACATAGATGACACC-3’) and cloned into the AscI site of the DsRED transformation vector. To generate the pS35:IQD22:mCherry translational fusion cDNA of IQD22 was amplified with primer IQD22 NcoI F (5’-CCATGGGAAAAGCGTCACGGTGGTTTA-3’ and IQD22 EcoRV R (5’-GATATCTCAGTACCTATACCCAATTGGCAT-3’), cloned in pJET2.1 and sequenced after which the fragment was cloned in pJD330 from which the GUS fragment had been removed. A MluI site which had been introduced by *in vitro* mutagenesis at the 3’ end of the IQD22 open reading frame was used to clone mCherry, which had been amplified with primers containing the MluI restriction site. The S35:IQD22:mCherry translational fusion was amplified with the same primers used for the pIQD22:GUS reporter, after which the fragment was blunted and cloned in BIN19. For the pIQD22:IQD22:mCherry construct the IQD22:mCherry fusion in pJD330 was amplified with IQD22 KpnI F5’-GGTACCCAACTTGCATTATTACTGTACCA-3’ and AscI nos and cloned into pJet2.1. The KpnI site was used to insert the IQD22 promoter fragment which had been amplified with KpnI F (5’-GGTACCCAACTTGCATTATTACTGTACCA-3’ and KpnI R (5’ -GGTACCTACTAATGAAAGTTACTTGACGAAT-3’). Clones where the promoter had inserted in the right orientation where isolated and the complete pIQD22:IQD22:mCherry product was amplified with (AscI F 5’-GGCGCGCCCTTTGCCAACGAATGTTAGTTGT-3’ and AscI nos 5’-GGCGCGCCATCTAGTAACATAGATGACACC-3’), cloned in pJet2.1, sequenced, excised with AscI, blunted and used for cloning in BIN19.

### Gas exchange measurements

Leaf gas-exchange and fluorescence measurements were performed simultaneously using a portable leaf gas exchange and fluorescence system (LI-6400XT and fluorescence system 6400-40; LI-COR, Lincoln, NE, USA). The measurements were performed on fully expanded leaves of 5 weeks old plants. Leaves were clamped on the 2 cm^2^ chamber of the LI-6400XT system and allowed to stabilize for 5 to 10 minutes at the following conditions: leaf temperature of 20°C, saturating quantum flux density of 1000 µmol m^-2^ s^-1^ (pre-tested); CO_2_ concentration in the cuvette of 400 µmol CO_2_ mol air^-1^.

Data for the CO_2_ response curves of photosynthesis (A/C^i^ curves) were collected at a range of ambient CO_2_ concentration (C^a^) in multiple steps: 400-300-200-100-400-450-550-700-1000-1200 µmol CO_2_ mol air^-1^. For every point, the output gas-exchange parameters (A, gs, E, among others) were recorded and light-adapted chlorophyll a maximum fluorescence (Fv’/Fm’) was measured after a flash of saturating light. The maximum carboxylation rate (Vcmax), electron transport rate (J) and mesophyll conductance were calculated from the A/Ci curves according to Ethier and Livingston (2004) using the A/Ci Curve Fitting 10.0 excel worksheet (available at landflux.org/tools). Chlorophyll content was measured using a SPAD meter (SPAD-502, Minolta; Japan) in the same leaves used for gas-exchange and fluorescence measurements. There were 5 replicates per treatment for all the measurements. Data was checked for normality and ANOVAs and *post-hoc* Tukey tests were performed for each parameter using the GenStat software.

Ethier G. J. & Livingston N.J. (2004) On the need to incorporate sensitivity to CO2 transfer conductance into Farquhar – von Caemmerer – Berry leaf photosynthesis model. Plant Cell and Environment 27(2):137–153. DOI: 10.1111/j.1365-3040.2004.01140.x

## Acknowledgements

The authors thank BBSRC (UK) for financial support under Institute Strategic Programme Grants BBS/E/C/000I0420. We would also like to thank Dr Joseph F McKenna (Oxford Brookes University) for the Arabidopsis 35S-TUA6-GFP seeds.

## Author Contributions

M.C.M conceived the project, M.C.M and S.K designed the experiments. M.C.M. conducted and analysed most of the experiments with assistance from S.K.; J.P.P. performed the measurement of photosynthetic parameters and analysed these data. M.C.M and S.K. wrote the manuscript with contributions from J.A.N. All authors commented on the manuscript before submission.

