## Supplemental Data for "The palisade-specific gene *IQD22* enhances photosynthetic capacity by phenocopying “sun leaf” architecture"

**Matthes et al IQD22**

**Supplemental Figures**

**Fig. S1**

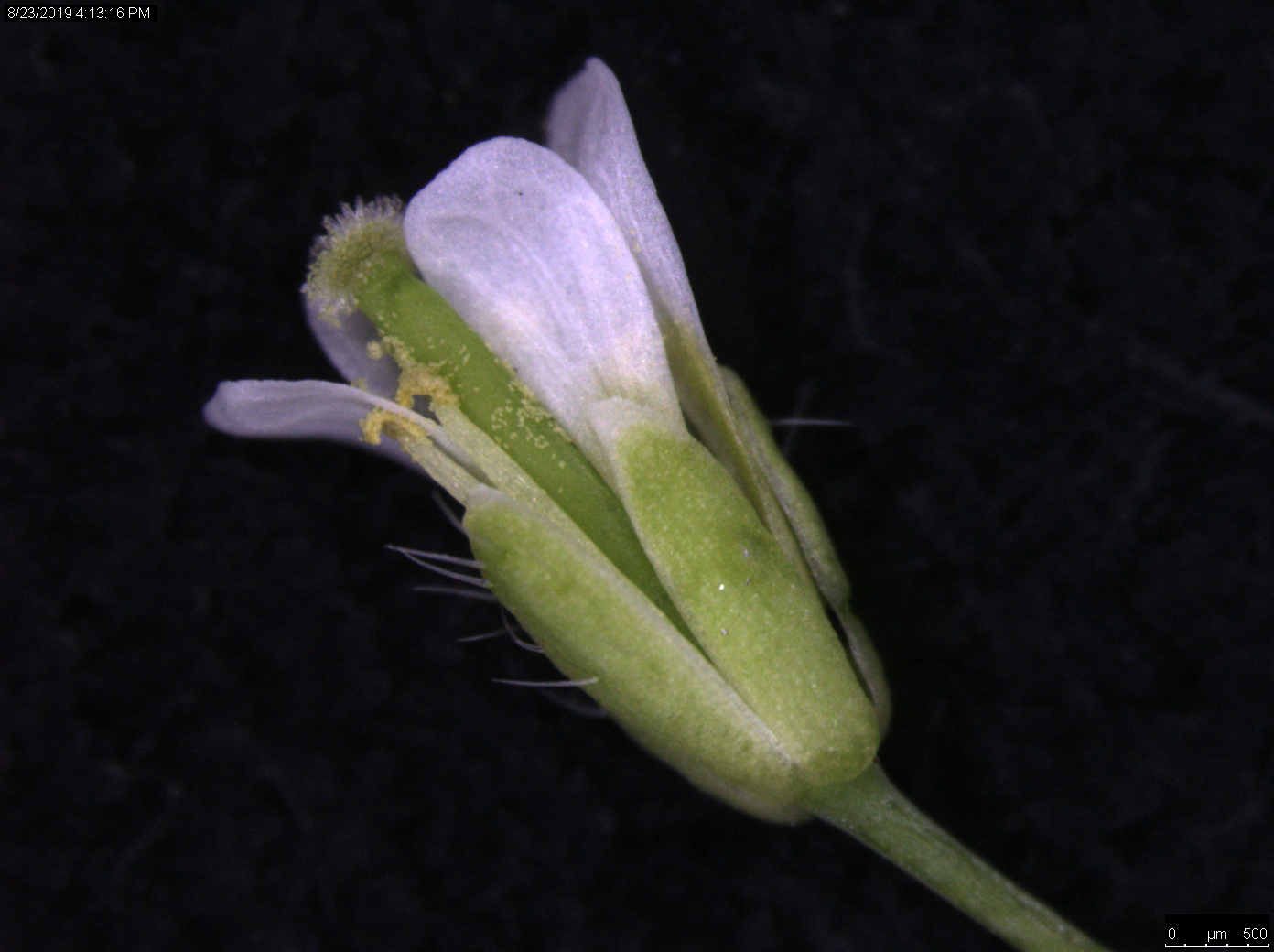

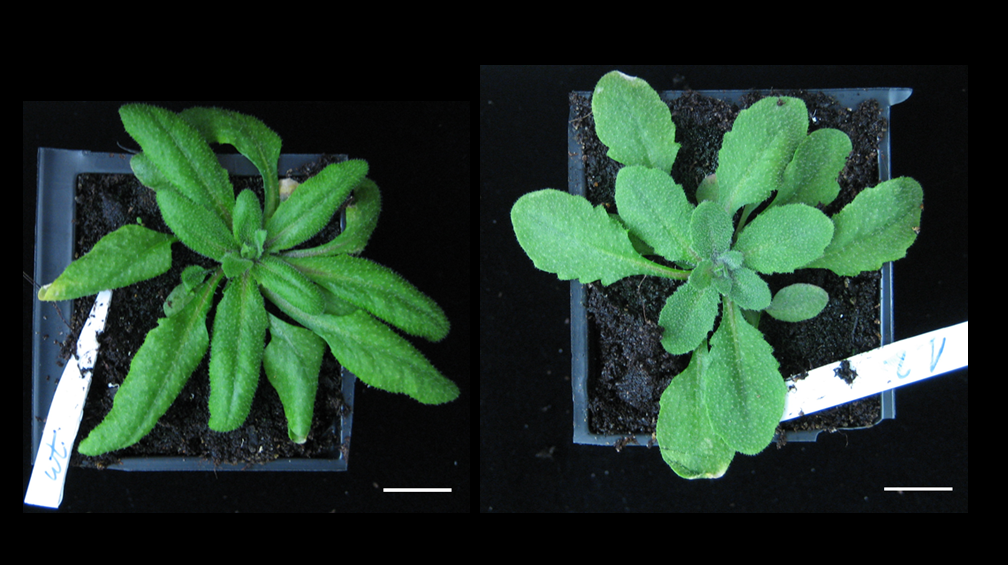

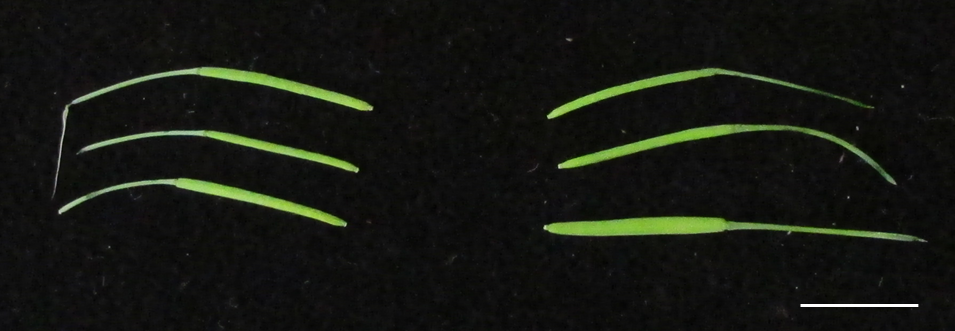

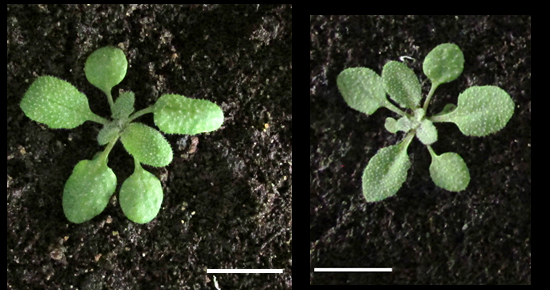

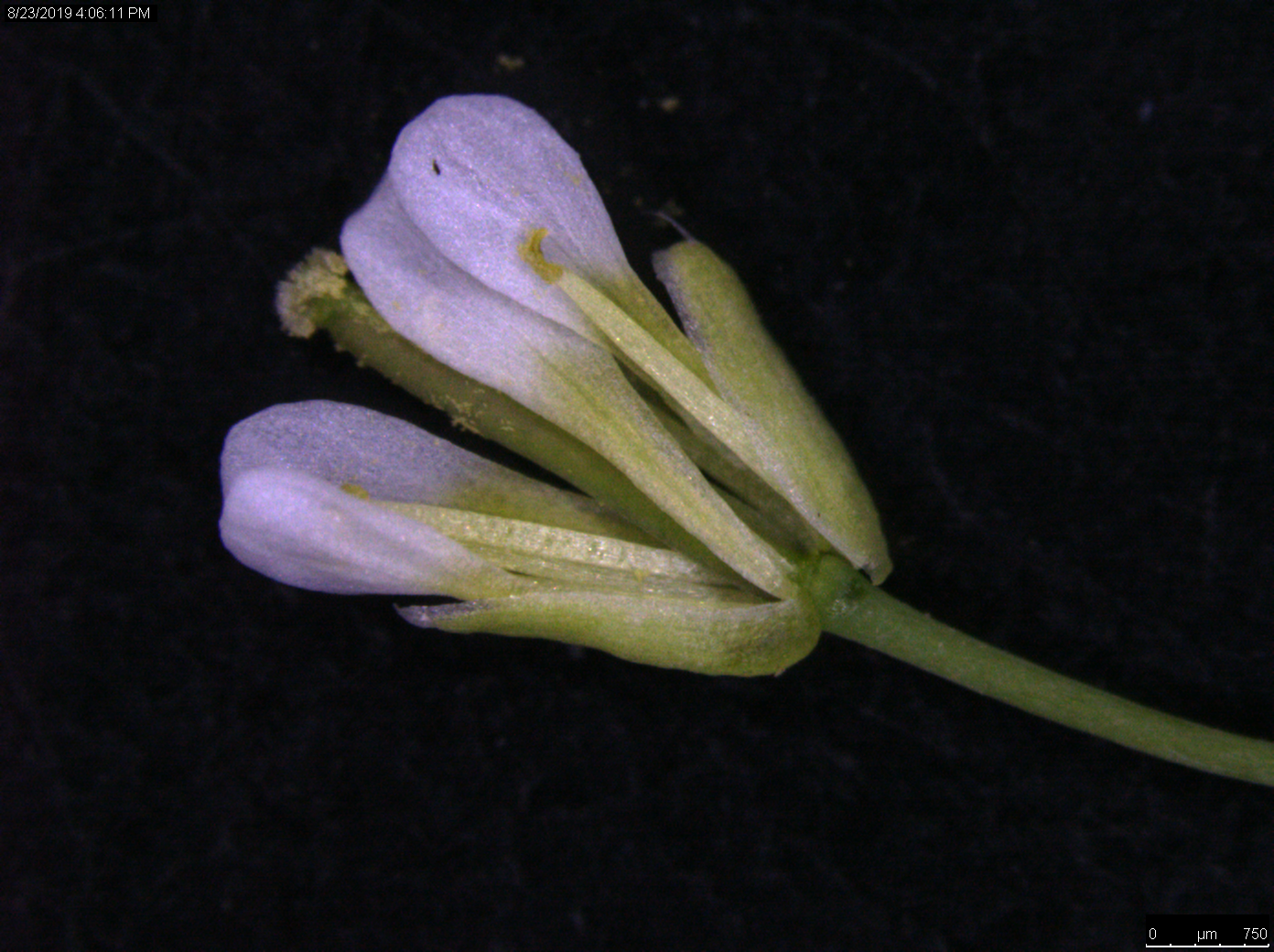

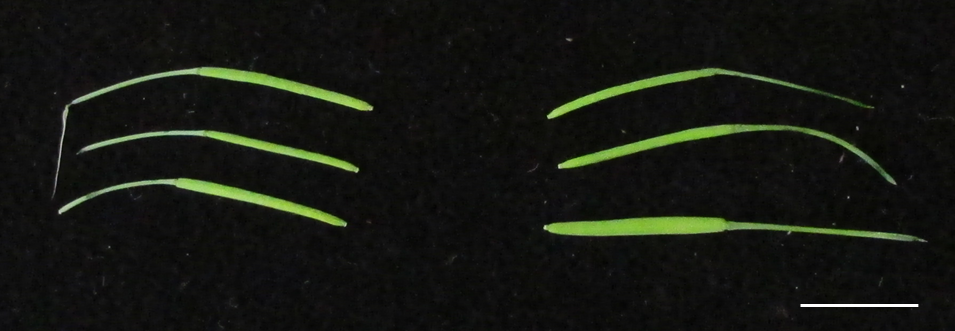

**a**

**b**

**c**

**d**

**Supplemental Figure S1:** Comparison of the phenotype of the WT and a representative transgenic pIQD22:IQD22:mCherry line at different developmental stages. **a**, 17 days old rosette and **b**, 25 days old rosette show that leaf shape in the transgenic line is altered whereas the flowers (**c**) and siliques (**d**) appear not to be affected by the expression of the transgene. Bar a, b = 1 cm ; c = 500 μm; d = 1 cm.

**Fig. S2**

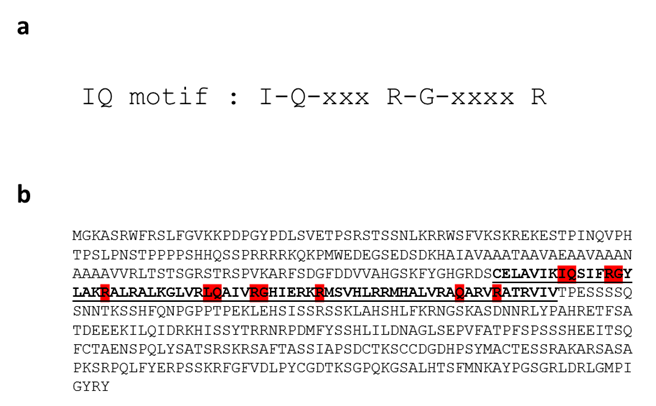

**Supplemental Figure S2: a,** Amino acid sequence of the signature motif of the IQ 67 domain present in 33 members of the *Arabidopsis thaliana* IQD family. The IQ67domain is characterised by the invariant spacing of up to three IQ motifs or of its more relaxed version [I/L/VQxxxRxxxxR/K]. **b,** Amino acid sequence of IQD22. The IQ67 domain is underlined and the conserved amino acids of the IQ motif are highlighted in red.

**Fig. S3**

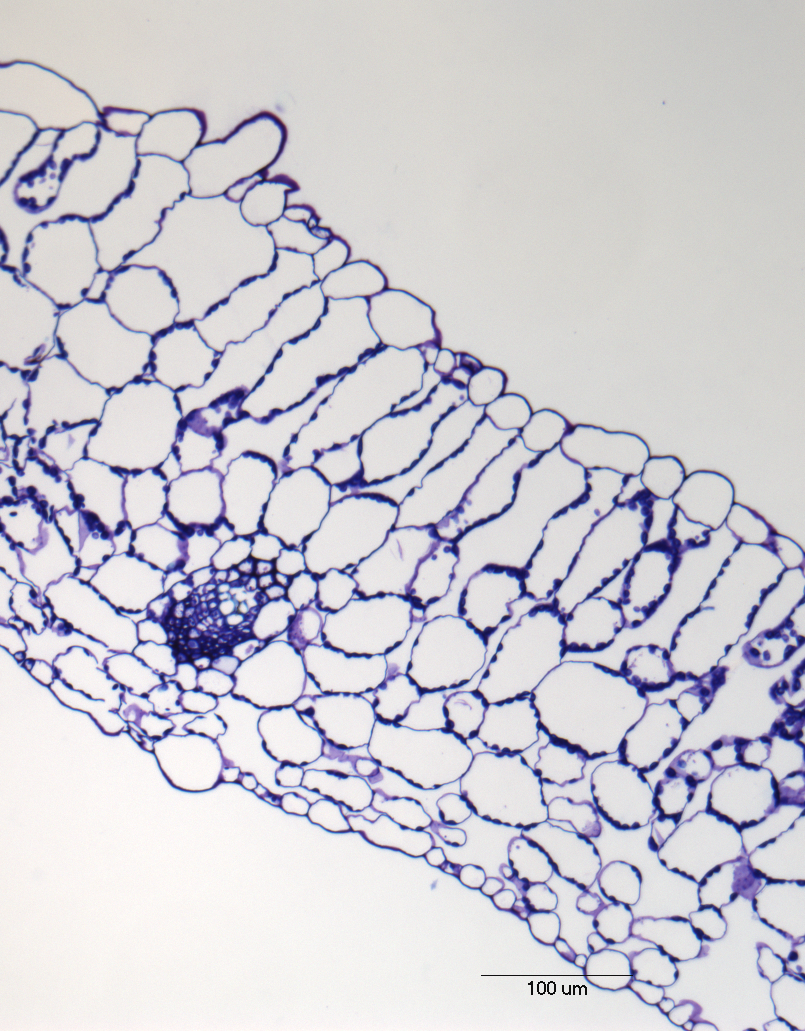

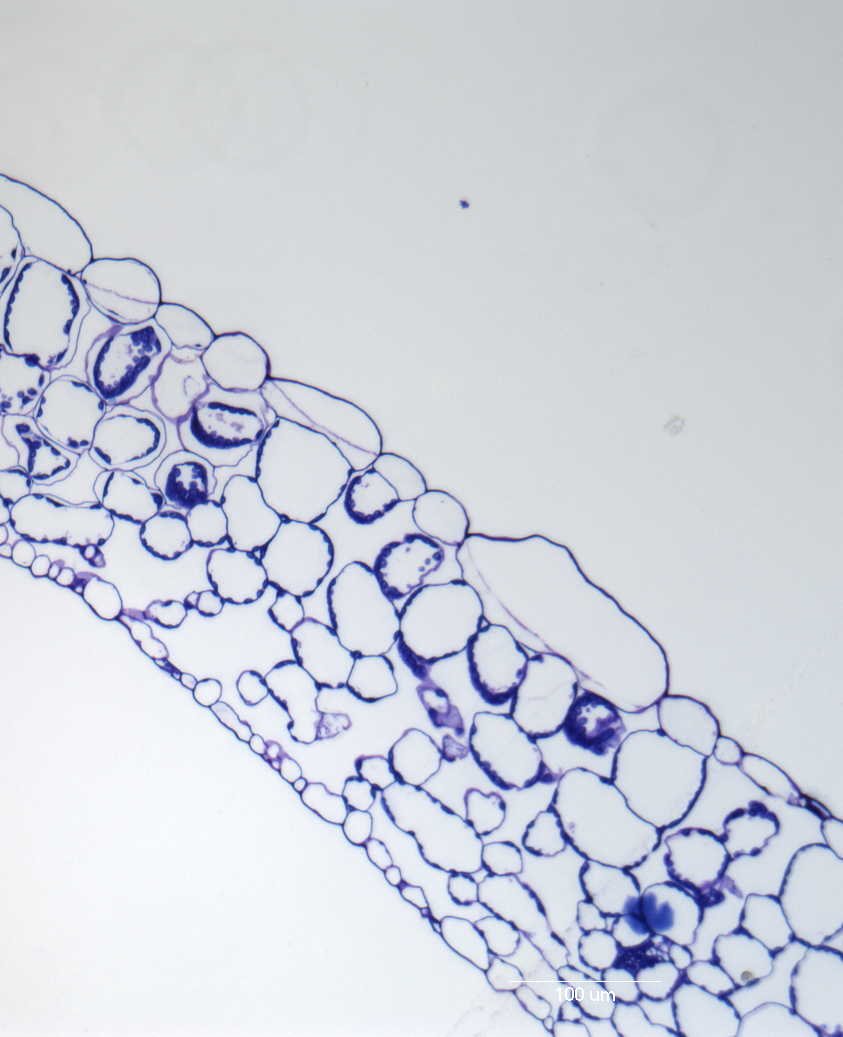

WT

pIQD22:IQD22:mCherry

Line 1

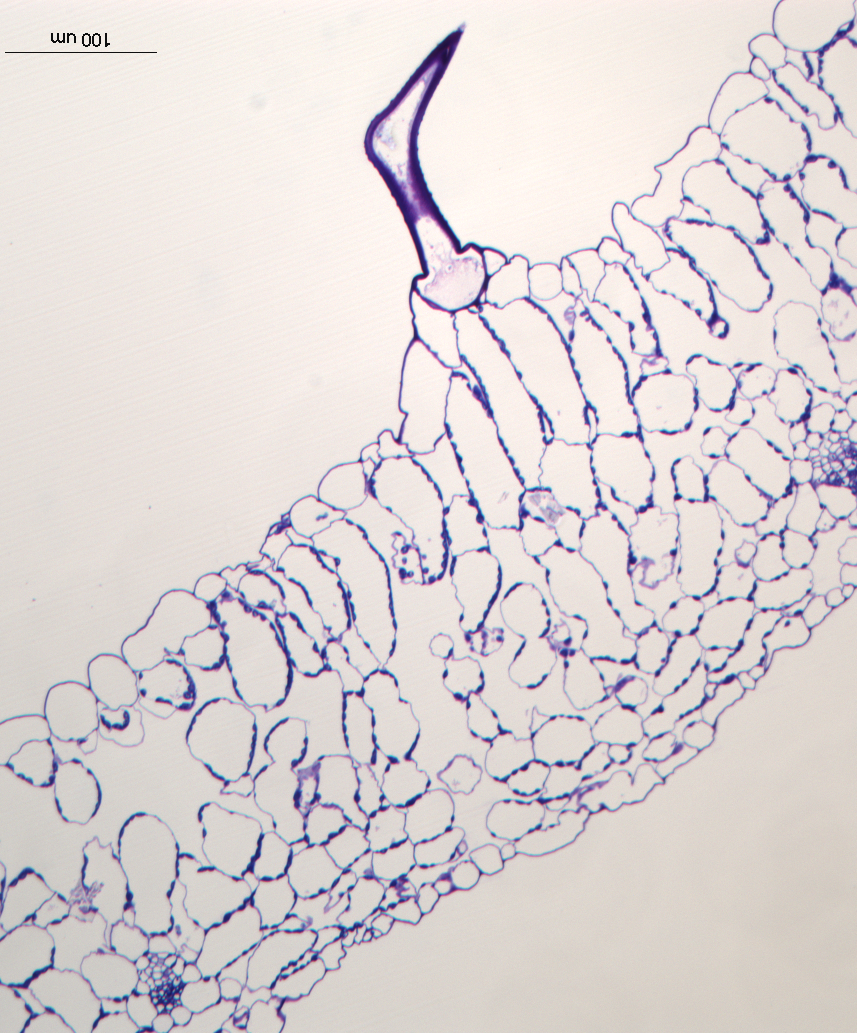

pIQD22:IQD22:mCherry

Line 2

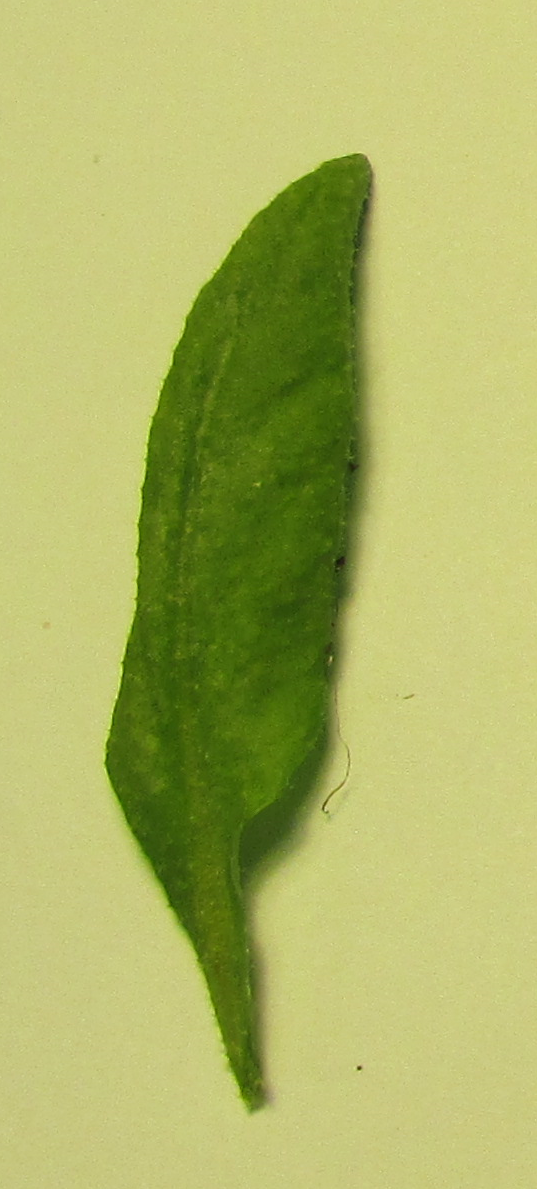

1

2

1

2

3

1

2

3

1

2

3

4

PM

SM

SM

PM

PM

SM

**Supplemental Figure S3: pIQD22:IQD22:mCherry expression leads to differentiation of additional palisade cell layers and elongation of cells within the tissue.** The same areas of mature leaves from 3 week old plants grown at 200 μmol m^-2^ s^-1^ white light were sampled. Cross sections of resin embedded samples were analysed by ImageJ for the generation of the data shown in Fig 1d. Numbers indicate the layers of palisade cells counted within the palisade tissue and show that in the transgenic lines palisade tissue has more layers. PM = palisade mesophyll, SM = spongy mesophyll. Bar = 100 μm.

**Fig. S4**

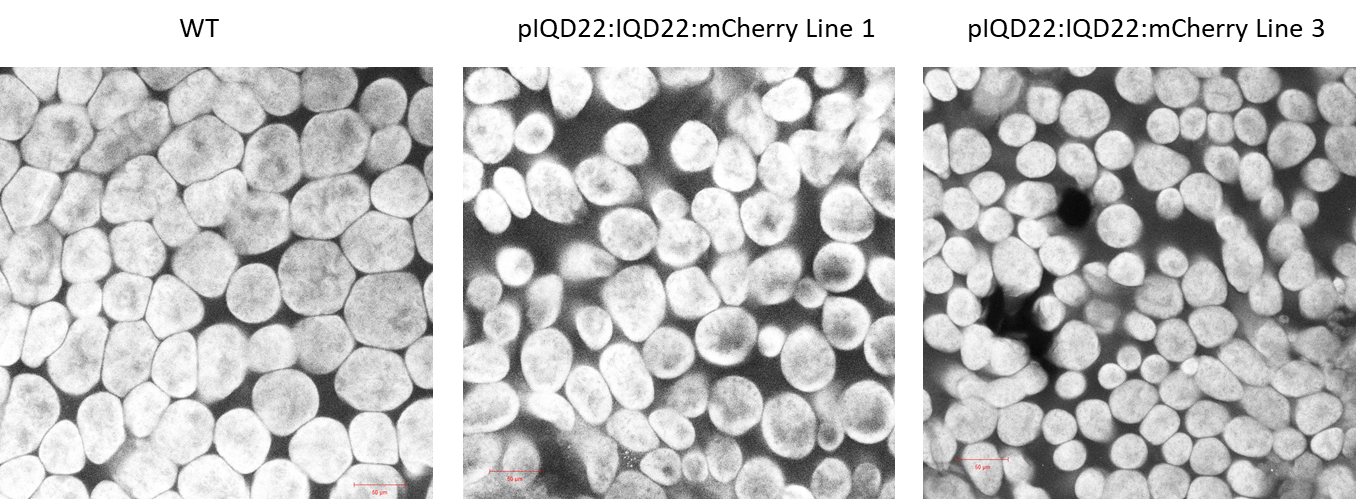

**Supplemental Figure S4: Palisade cell arrangement in the paradermal plane.** Fully mature leaves of WT and two independent transgenic lines were fixed and cleared with 70% ethanol for 24 h and subsequently in 100% lactic acid which was also used as mounting agent for microscopy. Leaf images were obtained with the Zeiss 780 LSM by imaging residual autofluorescence of chlorophyll (647-721 nm) after excitation with the 633 nm laser. Bar = 50 μm.

**Fig. S5**

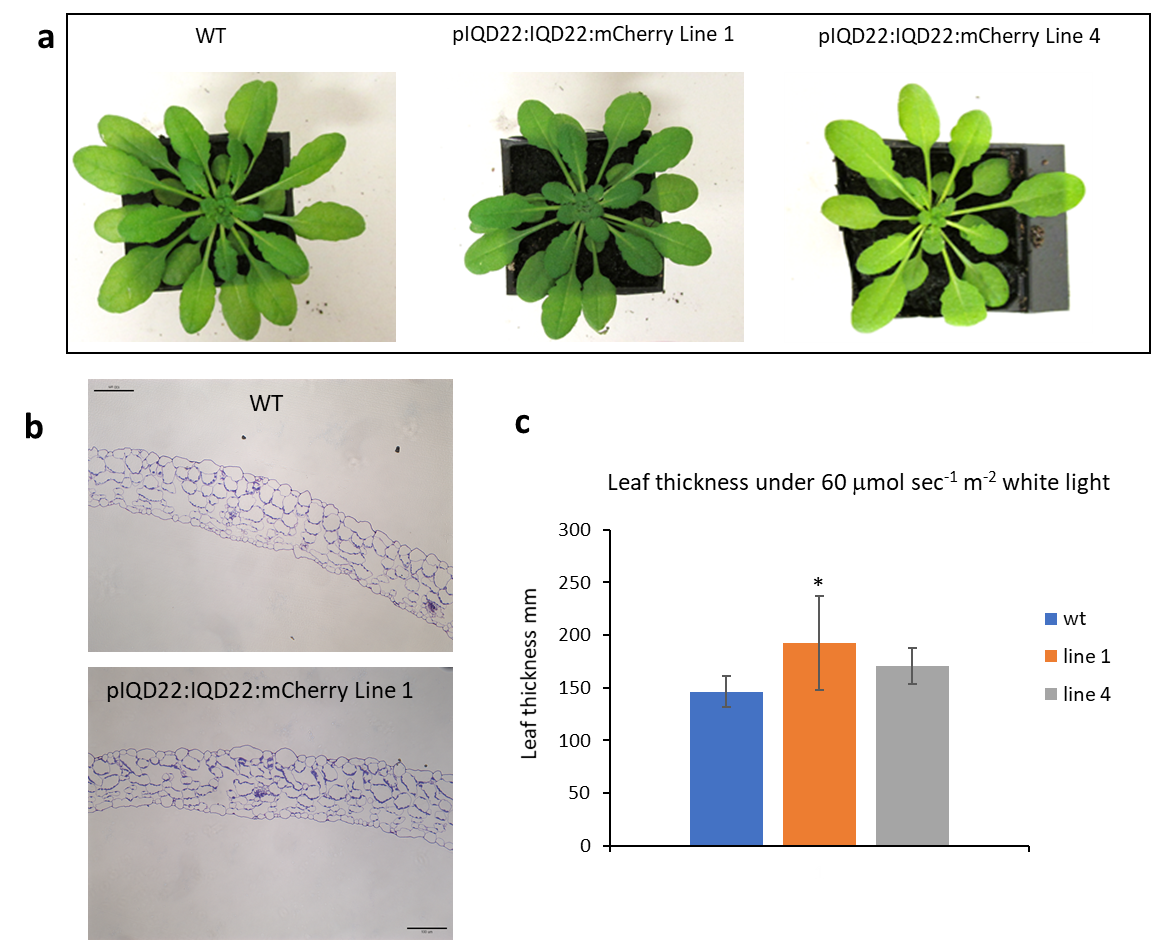

**Supplemental Figure S5: Development of increased leaf thickness in pIQD22:IQD22:mCherry transgenic lines is largely dependent on light intensities. a,** phenotype of 7 week old plants grown at 60 μmol m^-2^ s^-1^ white light shows no difference between the different genotypes. **b,** resin embedded cross-sections of WT and pIQD22:IQD22:mCherry line 1 show that the remarkable elongation of palisade cells as observed under 200 μmol m^-2^ s^-1^ white light does not occur under lower light intensities which is reflected in **c,** the measurement of leaf thickness. There is a slight but significant increase in leaf thickness in Line 1 compared to WT (t-test, p=0.01) whereas line 4 shows no significant increase. Bar = 100 μm.

**Fig. S6**

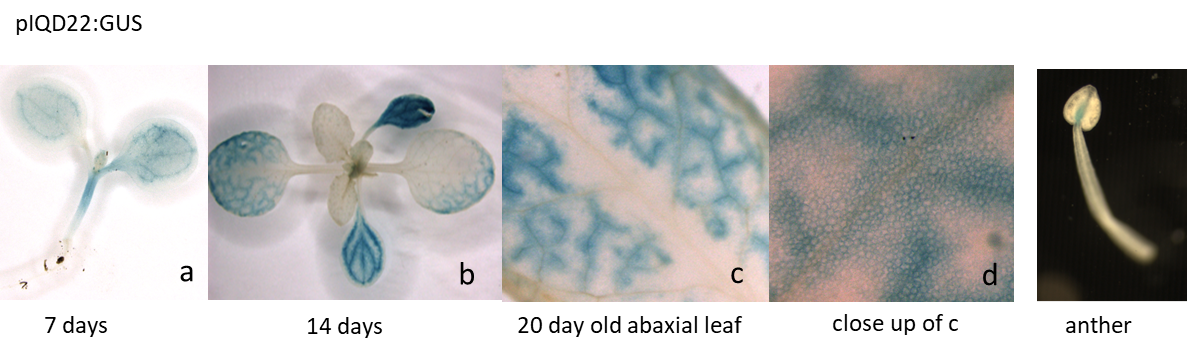

**Supplemental Figure S6: Expression of IQD22 as detected by pIQD22:GUS fusion.** Four independent lines containing the IQD22 promoter: GUS reporter fusion were grown in compost and cultivated under 200 μmol m^-2^ s^-1^ white light and long day conditions. Histochemical staining is shown for two independent lines at different developmental stages.

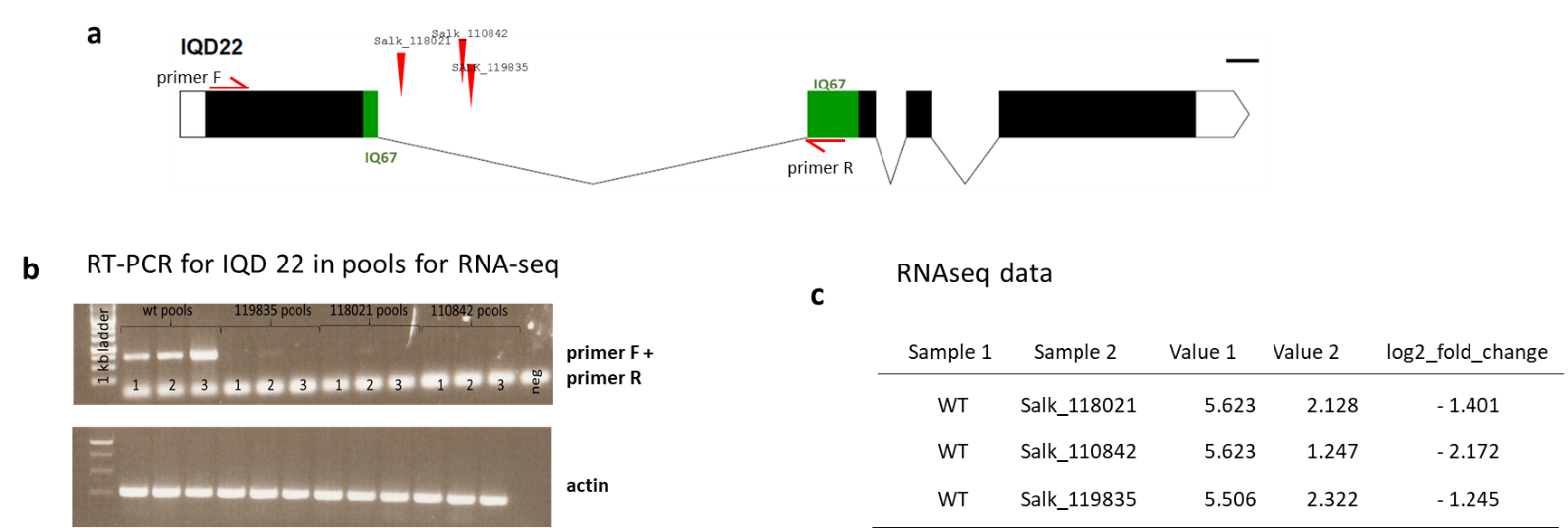
**Fig. S7**

**Supplemental Figure S7: a,** Scheme of the genomic organisation of IQD22 with the T-DNA insertion sites of the different lines analysed are indicated. (Primer F : 5’-CCATGGGAAAAGCGTCACGGTGGTTTA-3’ and primer R: 5’ TTGGAAGTGAGAAGATTTGGT-3’ show the localisation of the primers used for RT-PCR). Bar = 100 bp. **b**, RT-PCR of samples used for RNA-Seq. Each pool consisted of leaf 8/9 from 5 individual 4 week old plants, 3 replicates per genotype were analysed. **c**, Although RT-PCR on the RNA pools (**b**) indicated that they were true KO lines, subsequent RNA-Seq showed that these SALK-T-DNA lines were in fact knock downs lines and no true KO line have been isolated.

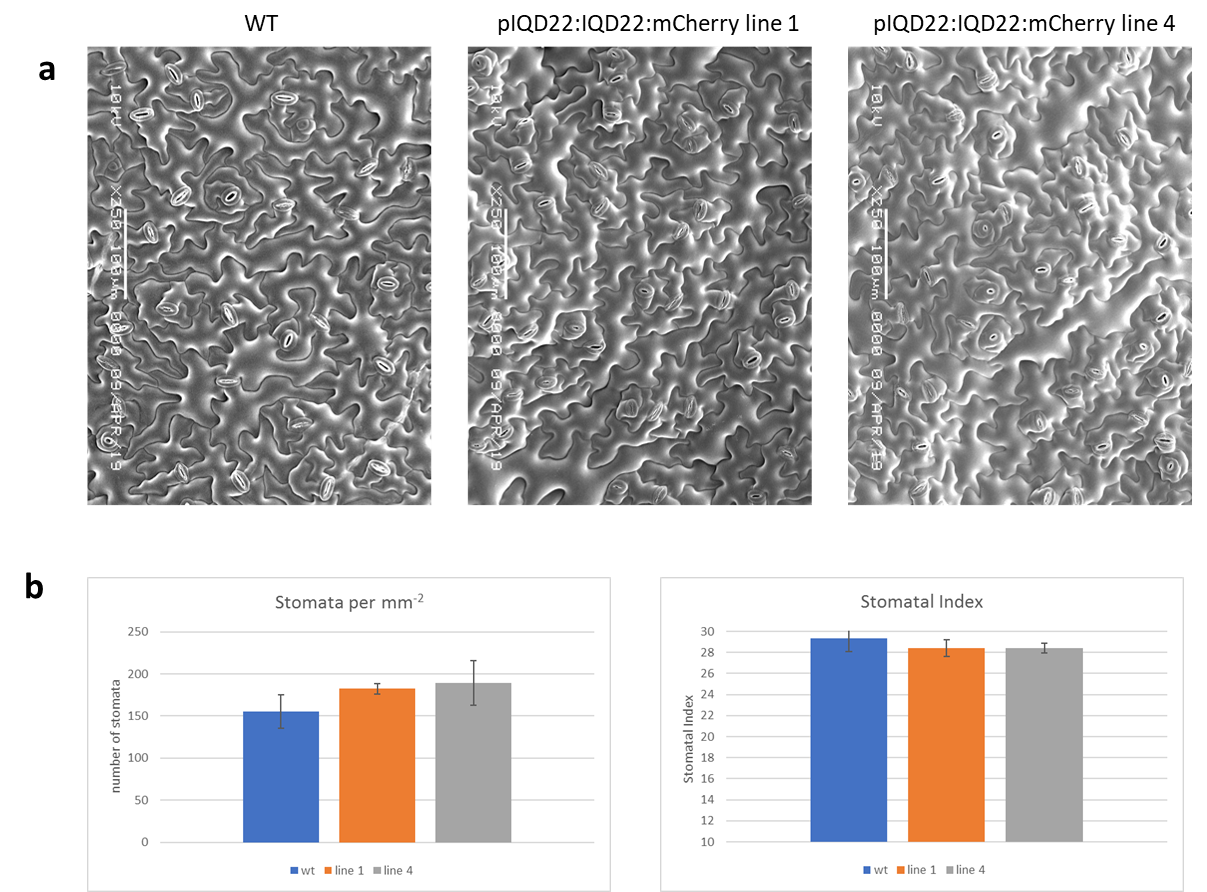
**Fig. S8**

**Supplemental Figure S8: Increase in photosynthesis is not the result of more stomata as stomatal index in WT and pIQD22_r_:IQD22:mCherry lines is not altered. a,** SEM images of the abaxial epidermis of mature leaves (leaf 8-9) of 45 days old plants used for counting of stomata. Bar = 100 μm. **b**, For determination of stomata per mm^-2^ and stomatal index, 3 replicates per line were imaged. For each replicate 3 different areas of the leaves were imaged and stomata and epidermal cells counted. T-Tests confirmed that no significant difference in stomata number and index occurred between the WT and transgenic lines.
